# A potential mechanism for the first-person inner sensation of memory provides evidence for a relationship between learning and LTP induction

**DOI:** 10.1101/085589

**Authors:** Kunjumon I. Vadakkan

## Abstract

Large number of correlations have been observed between behavioral markers of memory and long-term potentiation (LTP). However, there are different non-correlated findings that need explanations. Examples include a) a delay of at least thirty seconds for LTP induction after stimulation that does not match with milliseconds of time required for associative learning, and b) the deficiency of the LTP mechanism for providing a structure-function mechanism for working memory. By viewing memories as first-person inner sensations, a derived mechanism can explain various features of LTP and its mismatched findings with that of normal learning.

## 1 Introduction

A large number of correlations between behavior associated with memory retrieval and long-term potentiation (LTP) have been observed at various levels (Morris et al., 1986; McNaughton et al., 1986; Castro et al., 1989; Moser et al., 1998; Malenka, and Nicoll, 1999; Whitlock et al., 2006; Volianskis et al., 2015). In fact, this correlation is a key observation that can guide to understand the cellular-level events during learning and memory retrieval. The source and routing of the potentials that trigger motor neurons concurrent with the actual mechanisms that induce memory are expected to provide the crucial information how the cue stimulus evokes memory. Since no cellular changes are observed during memory retrieval, the memory likely results from a passive reactivation of the changes occurred at the time of learning. In this context, the learning-induced cellular mechanism and its maintenance are expected to have similarities with the cellular-level changes during LTP induction and its persistence. Understanding this is of paramount importance in filling the explanatory gap between the experimental findings in biochemical, cellular and electrophysiological fields (Barnes, 1995; Stevens, 1998; Goldberg et al., 2002; Lisman et al., 2003; McEachern and Shaw, 2005; Abbas et al., 2015) and their correlation with that of the motor changes occurring via speech and behavior that are being studied by the field of psychology (Shors and Matzel, 1997). By asking the question "What are the real conditions that the solution must satisfy?", the present work has examined results from a large number of LTP-related studies and provided evidence for an interconnecting cellular mechanism.

### 1.1 Long-term potentiation

Donald Hebb postulated that when an axon of a neuron is near enough to excite another neuron repeatedly, some growth process or metabolic change takes place in one or both of these neurons such that the efficiency of a neuron to fire the neighboring neuron increases (Hebb, 1949). This marked the beginning of the thought process towards examining the cellular-level changes that occur during learning. Hebb’s postulates describe synaptic changes that can facilitate future use of the same synapses during learning. In the following years, a patient named H.M who underwent removal of both hippocampi for treating intractable seizures suffered severe memory loss following the surgery. H.M was examined for memory loss by testing behavior and speech. H.M failed to show signs of memory retrieval for the events or items learned during certain period of time prior to the surgery and was unable to learn anything new (Scoville and Milner, 1957). This indicated that the hippocampi are involved in the storage and/or retrieval of memory. Following these findings, laboratory experiments were focused on the hippocampal tissue with the hope that any electrical event that can persist for long period of time can become a suitable marker for the increased synaptic efficiency in accordance with the Hebb’s postulates. Such an electrical change named LTP was observed in the hippocampal sections (Lømo, 1966; Bliss and Lømo, 1973).

The experimental steps of LTP can be described as follows. The hippocampus is removed from the animals and slices are prepared by retaining the connectivity between the different orders of neurons (Fig. 1) and are maintained at near-physiological conditions. When an electrode is used to stimulate a large number of recurrent Schaffer collaterals of the excitatory CA3 (Cornu Ammonis layer 3) layer neurons whose (presynaptic) terminals synapse with the dendritic spines (spines or postsynaptic terminals or postsynapses) (these terms are used interchangeably as follows: in the context of a synapse, postsynaptic terminal is used; in the context of a neuron, spine is used) of the neurons of the CA1 layer, a recording electrode placed extracellularly (for recording field potentials) at the main dendritic stem area of the CA1 neurons or intracellularly by patch-clamping one CA1 neuronal soma records electrical changes (for simplicity, mainly patch-clamping results are explained hereafter). Based on Hebb’s postulates, these electrical changes are expected to be a function of changes at the CA3-CA1 synapses. It was found that if a brief repetitive stimulation is applied initially at the Schaffer collaterals, then the application of a regular stimulus at the same location is sufficient to produce a potentiated effect (125 to 300% increase in the field excitatory postsynaptic potential (field EPSP)) (interpretation from Abbas et al., 2015) following a delay period of nearly 30 seconds (Gustafsson and Wigstrm, 1990) and even more than a minute.

**Figure 1:**
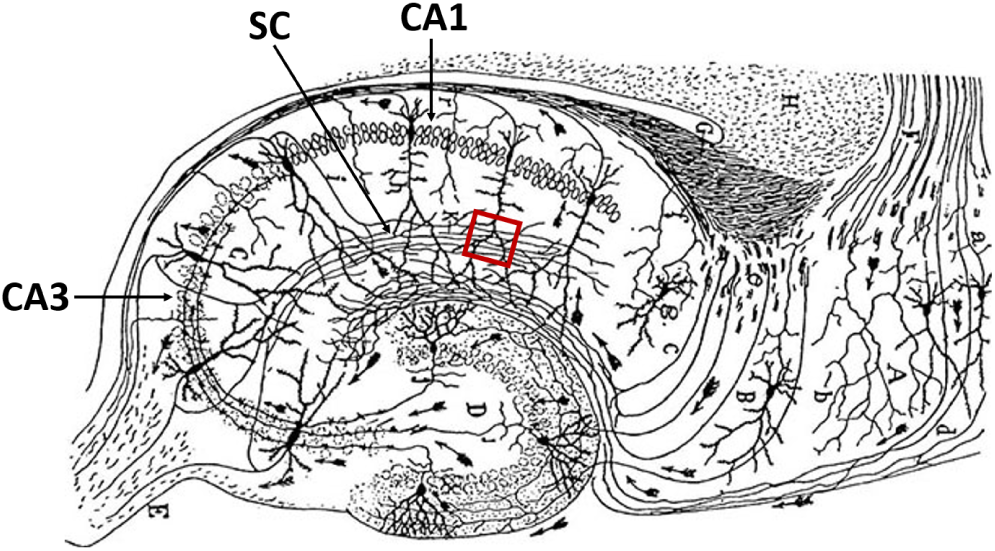
Diagram showing major hippocampal pathways: Schaffer recurrent collaterals (SC) connect between the CA3 and CA1 neuronal orders. Note that Schaffer collaterals cross at the middle of the dendritic tree of the CA1 neurons (approximately 100-250 micrometers from soma (Andrasfalvy and Magee, 2001). Red square: Shows a location where parallel Schaffer collateral axonal region at which their terminals synapse with the dendritic spines (postsynaptic terminals) of one CA1 neuron is drawn. Figure modified from (Cajal, 1909).

LTP induction requires a high amount of energy, which is provided by high frequency stimulation. A similar effect can be produced by using a single burst of strong activation (Remy and Spruston, 2007; Rose and Dunwiddie, 1986). By keeping the experimental preparation viable, an increased current arrives at the recording electrode in response to a regular stimulus applied at the same location of stimulation for several hours and even up to twenty-four hours. Following the initial observation of this finding, several experimental studies have found a correlation between the surrogate behavioral markers indicative of memory retrieval and LTP (Levy and Steward, 1979; Teyler and Discenna, 1984). Since then, synaptic level changes that can increase synaptic efficiency in accordance with the Hebb’s postulates were the focus of most investigations. Though initially discovered in the hippocampus, later experiments demonstrated LTP in different brain regions that receive different sensory inputs. LTP can also be induced at non-glutamatergic (Brown and McAfee, 1982) and inhibitory synapses (Ouardouz and Sastry, 2000). Several properties were attributed to LTP for considering it as an experimental correlate of the cellular mechanism for learning and memory (McNaughton et al., 1978; Levy and Steward, 1979; Andersen et al., 1977).

### 1.2 Models related to Hebbian plasticity

Several modified mechanisms were proposed to explain synaptic changes expected from Hebb’s postulates. These include: a) *Synaptic tagging hypothesis:* Possible intracellular mechanisms within single neurons were searched by conducting focussed electrophysiological studies. Normally, a single train of high frequency stimulation over the axonal bundles whose terminals synapse with a set of dendritic spines of a neuron elicits only an early-LTP. However, when this is preceded by stimulating a different set of synapses at the same neuron’s dendritic tree by using 3 or 4 trains of high-frequency stimulations (supra-threshold stimulations), late-LTP was recorded at the previous site where early-LTP was recorded. A demonstration of these findings was carried out in the hippocampus (Frey and Morris, 1997) and also by using cultured Aplysia neurons (Martin et al., 1997). This was proposed to occur by the generation of specific transient local synaptic tags at the synapses activated by a single train of stimulation that is expected to capture the gene products synthesized in response to the stimulation by 3 or 4 trains of high frequency stimulation. However, the time taken for the expression of genes fails to match with the physiological time-scales at which learning-induced changes are occurring. b) *Clustered plasticity and synaptic tagging model:* According to this model, bidirectional synaptic weight changes among the synapses at the spines of a dendritic branch are achieved through local translational enhancement (Govindarajan et al., 2006). This model incorporates synaptic tagging and capture as a mechanism for long-term memory where the dendritic branch is viewed as the location of operation of the integrative units (Govindarajan et al., 2011). Time-scale mismatch described for the synaptic tagging hypothesis also exists for the protein synthesis-dependent mechanism explained in this model. c) *Modified clustered plasticity model:* This view proposes that LTP at the synaptic region on the distal dendrites of the hippocampal CA1 pyramidal neurons requires cooperative synaptic inputs (Stuart and Spruston, 2015). Dendritic integration occurring at the level of the dendritic branches of a neuron whose still-unknown role in information processing is the main expectation of this model. All the above models associated with interaction between the spines of a single neuron do not provide a) mechanism for inducing inner sensations of different higher brain functions, and b) mechanism for providing two separate motor outputs from a single neuron each of which is part of two different associatively-learned stimuli.

### 1.3 Minimum requirements for an explanation

If the mechanism that induces LTP is same as that of learning, then there are different questions that require answers. The most basic questions include 1) What mechanism is taking place during learning that can be used to induce inner sensation of memory at the time of its retrieval and can also be observed during LTP induction? 2) What molecular events taking place at physiological time-scales during learning match with that taking place during LTP induction? Since it was only possible to correlate LTP with the surrogate behavioral markers of memory retrieval, the above questions cannot be answered directly. However, it provides an opportunity to make a hypothesis for a mechanism of induction of inner sensation of memory concurrent with the surrogate makers of memory retrieval using a mechanism taking place at the time of associative learning and test whether matching molecular events occur during LTP induction. The latter can be experimentally verified. During normal learning, only existing receptors or that can be inserted at the postsynaptic membrane within the normal time-scales will be used. Therefore, it is required to examine the molecular events taking place at comparable time-scales during LTP induction. Since retrieval-efficient learning can take place within milliseconds of time and since there is a delay of at least 30 seconds (Gustafsson and Wigstrm, 1990) to one minute for the induction of LTP after the high-energy stimulation, how can the latter be explained by a mechanism occurring during learning?

The results of the LTP experiments have been interpreted in terms of an increase in the synaptic efficiency in accordance with Hebb’s postulates. Even though increasing the synaptic efficiency may facilitate future use of the same synapses along the stimulated pathways, it is short of providing a learning-induced change from which memory of one item can be induced at the arrival of the second stimulus at the time of memory retrieval. The actual mechanism is expected to answer the question "How can the changes at the synapses through which the cue stimulus propagates induce the memory of the associatively-learned second stimulus?" In order to understand the exact mechanism, it is necessary to view memories in their exact nature as first-person inner sensations occurring concurrent with the behavioral changes and then address the question "What type of a change should occur between two associatively-learned stimuli during learning that will allow one of the stimuli (cue stimulus) to induce the inner sensation of memory of the second stimulus concurrent with behavioral motor actions reminiscent of the second stimulus?" This necessitates learning-induced changes to occur at the locations of convergence of sensory stimuli. Since no cellular changes were noticed at the time of memory retrieval, a passive reactivation of learning-induced change is expected to occur at the time of memory retrieval. Since the inner sensations of different memories are similar in nature, maintenance of learning-induced change occurring for varying periods of time is expected to explain different memories classified based on their duration of persistence. The maintenance of learning-induced changes and persistence of LTP for different periods of time and their capability to reverse back to the ground state have similarities. These increase the expectations for finding the matching cellular-level changes occurring during learning and LTP induction.

Several gene expression changes, the synthesis of new proteins and their modifications have been found related to both behavioral markers of memory retrieval and LTP (Herring and Nicoll, 2016; Madison et al., 1991), even though they occur more slowly than the physiological time-scales at which learning and memory retrieval take place. Since memory can be retrieved instantaneously following learning without having to wait for the biochemical sequence of events observed during and after LTP induction to take place, it leads the question "What cellular change occurring at physiological time-scales during learning is sufficient to lead to the induction of first-person inner sensation of retrieved memory?" It is necessary to separate the slowly occurring learning-induced cellular changes from those that are critical for learning and for inducing memories at physiological time-scales (Abbas et al., 2015). A flow chart diagram of the expected sequence of steps necessary to arrive at the explanation is given in Fig. 2.

**Figure 2:**
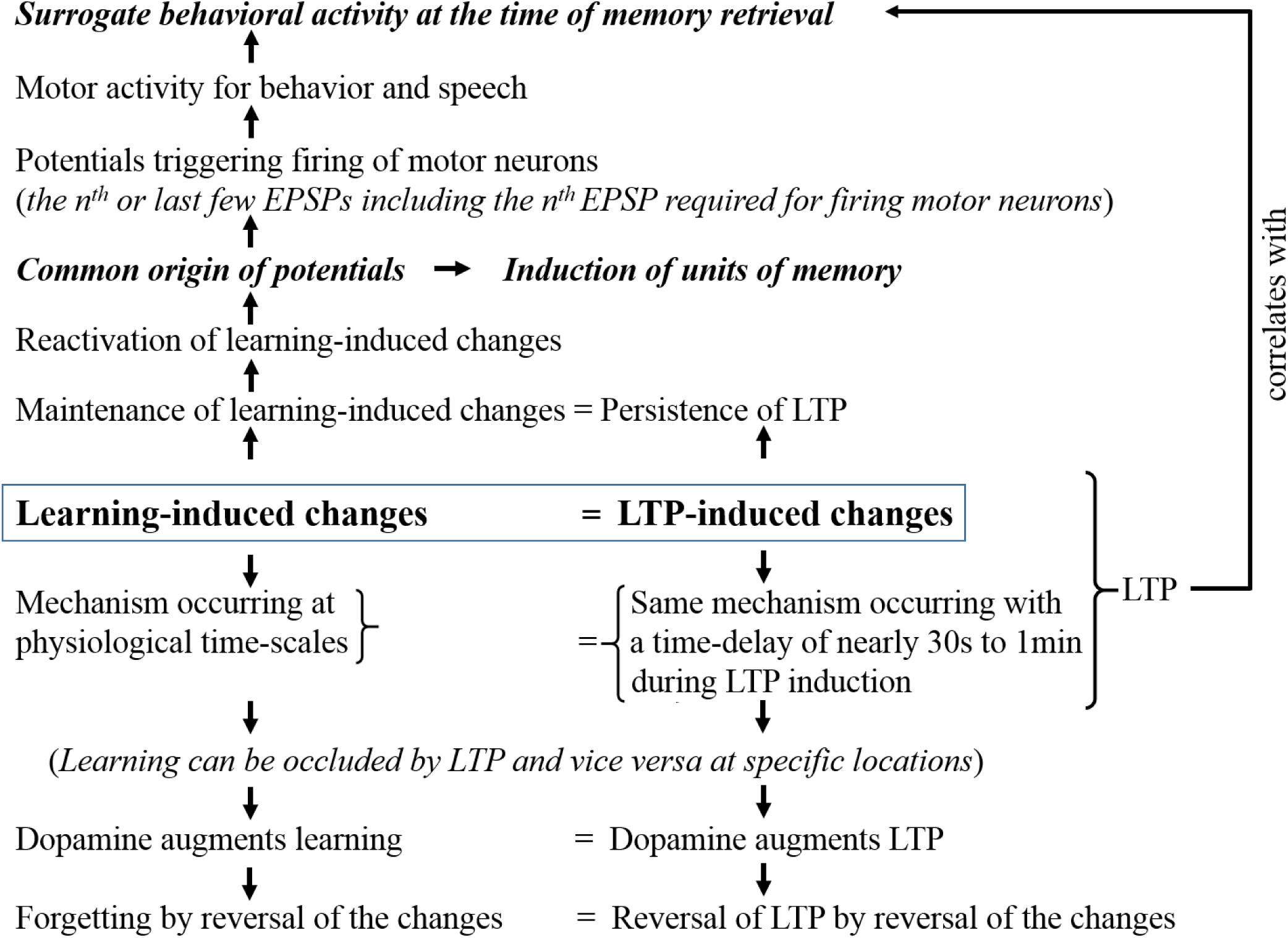
Steps that are necessary to prove the correlation between associative learning and LTP induction. These steps begin by searching for the source of potentials required for behavioral motor actions occurring concurrently with the induction of inner sensation of retrieved memories. Since cellular changes are absent during memory retrieval, a mechanism for the passive reactivation of a learning-induced changes is expected to occur. The learning-induced cellular change is also expected to correlate with LTP-induced changes occurring following a delay.

## 2 Derivation

### 2.1 Mechanism during learning from which memory can be induced

For learning-induced changes to take place at the locations of convergence of two associatively-learned stimuli, the inputs arriving through their presynaptic terminals must undergo a functional convergence at some point. Since there are only synaptic regions between the stimulating and recording electrodes in LTP experiments, the cellular mechanism of LTP is expected to take place at the level of the synapses - either at the synapses or between the synapses. This also indicates that the correlation of LTP with behavioral motor actions is likely taking place through a synapse-mediated mechanism. What synaptic level changes can a) induce inner sensations of memory, b) provide additional potentials for behavioral motor action, and c) explain LTP-induced changes? From a learning perspective, this can be examined as follows. For the cue stimulus to provide the source of potentials for both the induction of inner sensation memory and concurrent motor action at the time of memory retrieval, a reactivatible interaction is expected to occur between the associatively-learned stimuli at the locations of their convergence at the time of learning. At the locations of convergence, the axonal terminals (presynaptic terminals) of the two pathways synapse with the dendritic spines (postsynaptic terminals). Even though the importance of dendritic spines in cognition and its disorders are known (Koch et al., 1992; Penzes et al., 2011; Gonzlez-Burgos, 2012), it is not known whether the converging presynaptic terminals synapse on to the dendritic spines of the same of postsynaptic neuron or not. If they synapse to the same neuron, both stimuli will lead to the activation (spike or firing or action potential) of this same neuron and the identity of the associatively-learned stimuli beyond this neuronal level cannot be maintained. Moreover, when a neuron receives inputs during either sub-threshold or supra-threshold activations, these inputs do not contribute towards the neuronal firing and may not provide expected specific outputs. These indicate that the converging presynaptic terminals are likely synapsing on to the dendritic spines of different neurons as a rule. Exceptions may be present.

The converging presynaptic terminals from two associatively-learned stimuli are expected to induce certain interactive changes at the level of their synapses during learning. These changes provide specific signatures such that at a later time when one of the stimuli arrives as a cue stimulus, it will induce the inner sensation of memory of the second stimulus. What is the ideal location between the synapses of the converging presynaptic terminals that the interaction must occur during learning? The result of the interaction must allow the cue stimulus to induce the first-person inner sensation of memory of the second stimulus and also provide potentials to activate higher order neurons that belong to the second stimulus to produce corresponding behavioural motor actions. Since neurotransmission is taking place unidirectionally, the activation of the postsynaptic terminal (dendritic spine) can be viewed as equivalent to the activation of a synapse. Hence, interaction (a link formation) between the spines that belong to different neurons (as derived from the previous paragraph) can be examined for a suitable mechanism. In this regard, an interaction between the readily LINKable (capitalized to denote its significance) spines called inter-postsynaptic functional LINK (IPL) occurring at physiological time-scales was found to be suitable to explain the learning-induced changes (Fig. 3A) (Vadakkan, 2013; 2016a). The finding that the mean inter-spine distance is even greater than the mean diameter of the spine heads of the pyramidal CA1 neurons (Konur et al., 2003) indicates that in ideal situations, IPL formation takes place between the spines that belong to different postsynaptic neurons. This derivation can be subjected to experimental verifications. This will allow maintaining the specific motor outputs related with each of the associatively-learned stimuli at the time of memory retrieval.

**Figure 3.**
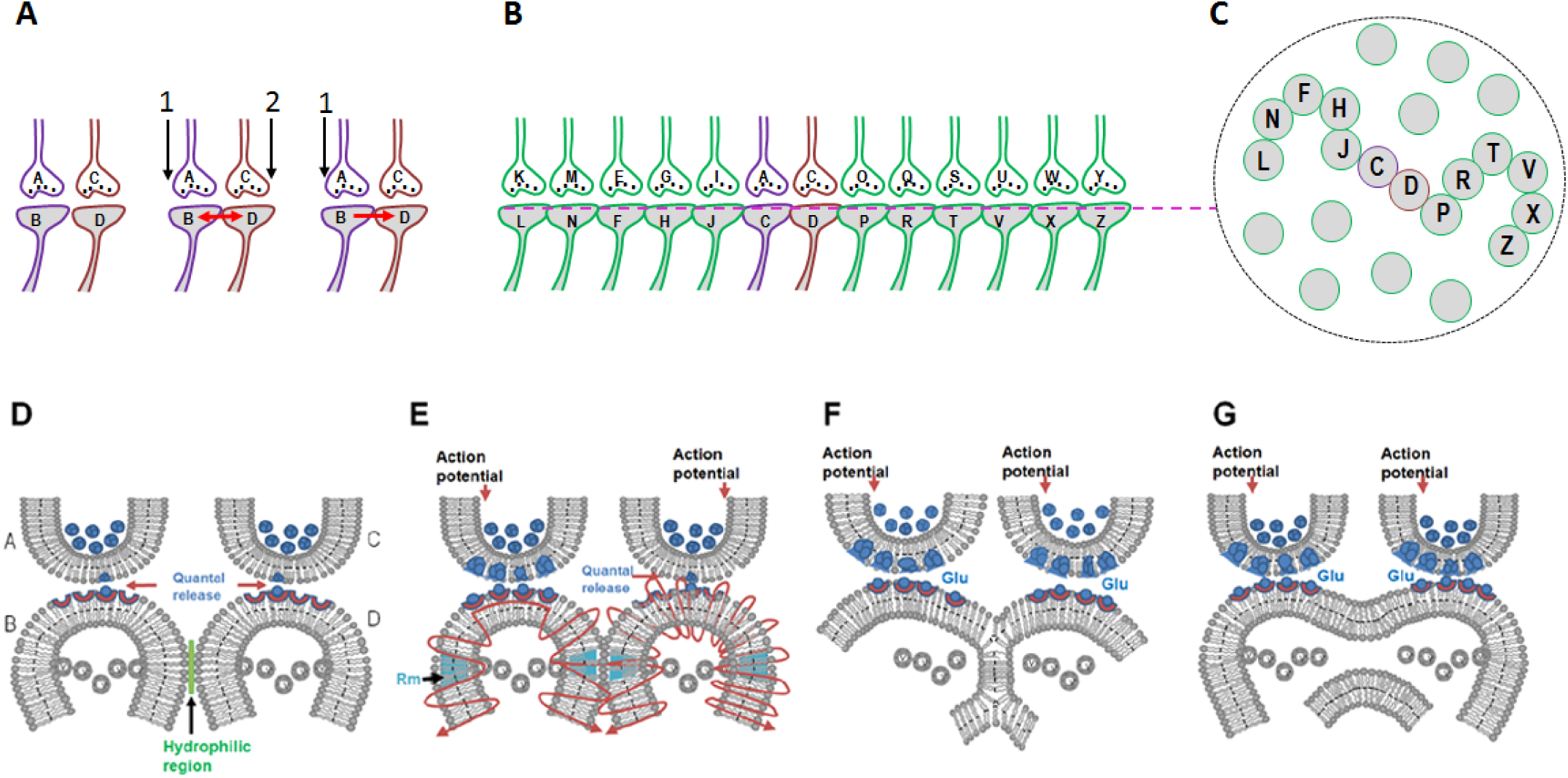
Inter-postsynaptic functional LINK (IPL) formation, clustering of inter-LINKed post-synaptic terminals (spines) and different types of IPLs. A) Left: Two synapses A-B and C-D are shown side by side. Note that their postsynaptic terminals B and D are abutted to each other. Middle: Simultaneous arrival of stimuli 1 and 2 at presynaptic terminals A and C during learning leads to the formation of an IPL between their postsynaptic terminals B and D. The formation of the IPL is a function of simultaneous activation of postsynaptic terminals B and D. Right: During memory retrieval in the presence of one of the stimuli (stimulus 1) the IPL is re-activated, resulting in the activation of postsynaptic membrane D. This results in the induction of semblance of activity arriving from presynaptic terminal C as an intrinsic systems property. The synaptic activities at synapses A-B and C-D are essential during learning and the synaptic activity at synapse A-B is essential for the reactivation of IPL B-D to induce the unit of inner sensation. B) Formation of IPLs with spines that have already made functional LINKs with other spines during prior learning events will lead to the clustering of functionally LINKed spines. The serially inter-LINKed spines L, N, F, H, J, C, D, P, R, T, V, X, and Z form an islet. Note that the inter-LINKed spines are arranged side-by-side. C) A cross-sectional view through the postsynaptic terminals within the islet of inter-LINKed spines shown in Figure B. Note the presence of eight other independent spines in the selected region of interest (D-G): Different types of IPLs. D) Presynaptic terminals A and C with synaptic vesicles inside (in blue color). The continuous quantal release is represented by one vesicle at the synaptic junction. Spines B and D have membrane-bound vesicles marked V containing AMPA receptor GluR1 subunits inside them. Spines are normally separated by a hydrophilic region between them (in green). Simultaneous activity arriving at the synapses leads to the enlargement of the spines and removal of the hydrophilic region between them that further leads to an electrically-connected close contact between them (not shown). Since it is a process requiring large amounts of energy, it is a rapidly reversible IPL and is responsible for working memory and short term potentiation (STP). E) Further enlargement of the spines and membrane reorganization (at the membrane segments marked Rm) secondary to AMPA receptor subunit vesicle exocytosis at the lateral borders of the spines can lead to reversible partial hemi-fusion between the spines. It is responsible for short-term memory and early LTP. F) Complete hemifusion between the spines that can be reversed and also stabilized. It is responsible for short- and long-term memories and LTP maintenance. G) Inter-spine fusion. Small areas of fusion may reverse back. Note that electrical continuity is maintained between the spines. Reversible fusion changes are expected to occur during LTP (Figures modified from Vadakkan, 2013; 2016a).

The mechanism for establishing an IPL should match with the properties of reversibility (that explains forgetting at different time periods after associative learning), stabilization (for long-term memory), augmented stabilization in certain conditions (e.g. motivation-promoted learning), and reactivation (for memory retrieval) (Vadakkan, 2016a). Continued associative learning events will inter-LINK additional spines with the initially inter-LINKed spines. This will generate groups of inter-LINKed spines called islets of inter-LINKed spines (Figs. 3B, C). A spectrum of molecular and cellular changes can explain IPL formation (Vadakkan, 2016a). These include close contact between the postsynaptic membranes by hydration exclusion and inter-postsynaptic membrane hemifusion (Figs. 3D–F). Since hemifusion is an intermediate stage of the membrane fusion (Fig. 3G), artificial stimulation conditions of LTP may show fusion changes that may reverse back or persist for different durations.

IPLs are expected to last for different periods of time. Reactivation of the IPL by the cue stimulus is expected to induce an inner sensation (semblance) for the associatively-learned second stimulus. Details of the derivation and mechanism of induction of semblances were explained previously (Vadakkan, 2013) and are summarized in Fig. 4. Brifley, it evolved from asking the question "What local and system conditions can induce the property of first-person inner sensations at the time of lateral entry of depolarization through the IPL towards the inter-LINKed spine?" Concurrent with the induction of inner sensations, potentials arriving at the inter-LINKed spine can lead to the activation of the latter’s neuron if it is held at a sub-threshold activated state. If this neuron is a motor neuron or can activate a motor neuron at its higher neuronal orders, then it can contribute towards the behavioral motor action corresponding to the retrieved memory. Inhibitory interneurons and feedback circuits can regulate the crossing of the threshold limits of these neurons.

**Figure 4.**
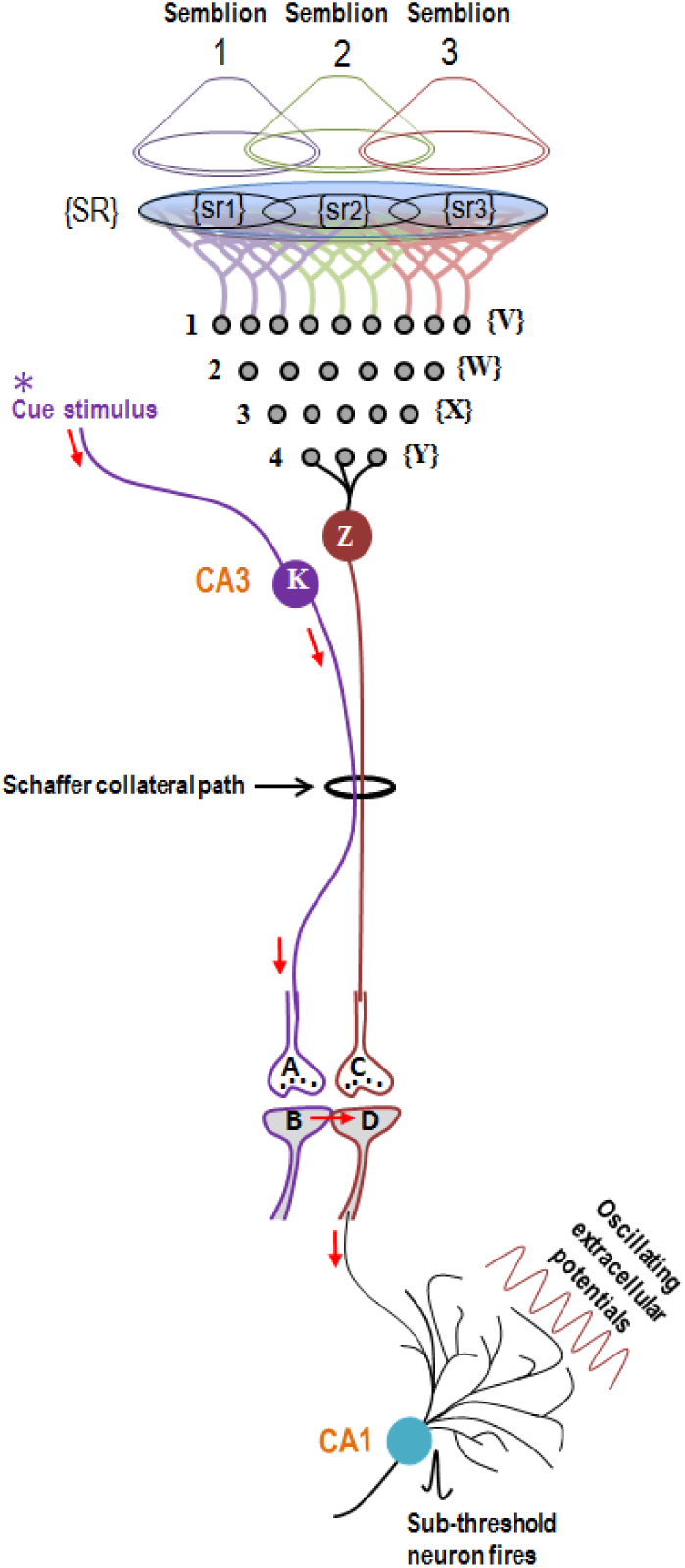
Induction of a unit of inner sensation and the concurrent firing of a sub-threshold neuron. The gray circles placed in rows and numbered from 1 to 4 are the different neuronal orders starting from the sensory receptor level. Neurons of the hippocampal CA3 layer (marked K and Z) are shown as neuronal order 5. When the cue stimulus reactivates IPL B-D, it activates postsynaptic terminal D from a lateral direction and induces a unit of inner sensation. In order to estimate the sensory identity of the inner sensation, a retrograde extrapolation from postsynaptic terminal D towards the sensory receptor level is undertaken. This retrograde examination constitutes the first-person approach. Postsynaptic terminal D is activated by CA3 neuron Z. Spatial summation of nearly 40 EPSPs (from nearly 40 dendritic spines out of the nearly tens of thousands of dendritic spines of each pyramidal neuron) or temporal summation of less than 40 EPSPs triggers an action potential at the CA3 neurons axon hillock (Normally, these numbers are likely to be higher since the EPSPs degrade as they arrive at the cell body). Let the set of all combinations (for the spatial summation of EPSPs) and permutations (for the temporal summation of EPSPs) of the neurons whose activity through normal synaptic transmission and depolarization order 3. By extrapolating arriving through the IPLs be Y in neuronal order 4. Neurons in set Y in turn receive inputs from dendritic spines and through the re-activation of IPLs from activity arriving from a set of neurons X in neuronal in a retrograde fashion towards the sensory receptor level, it is possible to determine the set of sensory receptors SR and from the latter the sensory identity of the cellular hallucination can be determined. In other words, dimensions of inner sensations resulting from the activation of postsynaptic terminal D through the re-activation of IPL B-D will be related to a sensory stimulus that can activate sensory receptors in set SR. It is likely that the activation of subsets of a minimum number of sensory receptors from set of SR, for example, sr1, sr2, and sr3 is sufficient to activate postsynaptic terminal D. A hypothetical packet of minimum sensory stimuli capable of activating one of the above subsets of sensory receptors that can activate postsynaptic terminal D is called a semblion. This is considered the basic unit of the inner sensation of memory. These sensory units have no orientation. The lateral activation of postsynaptic terminal D can induce a large number of semblions (Figure modified from (Vadakkan, 2013).

### 2.2 Suitability of the IPLs in explaining LTP

During LTP induction, the high-energy stimulation of a group of axonal terminals induces certain changes that need to be discovered. Following a delay period of nearly 30 seconds (Gustafsson and Wigstrm, 1990) and even more than a minute, a regular stimulus applied at the same location induces a potentiated effect of up to 300% increase in the field EPSP (interpretation from Abbas et al., 2015). Since it was found that local synaptic depolarization and/or dendritic spikes can mediate a stronger form of LTP than alternative methods for its induction (Hardie and Spruston, 2009), the present work examines this direct mechanism to find its correlate at the time of learning. Only a fixed number of spines of the postsynaptic CA1 neuron (from whose soma the recording is carried out) synapses with a fixed number of axonal terminals (presynaptic terminals) of the Schaffer collaterals. A change occurring at these fixed number of synapses alone to produce a potentiated effect of up to 300% following an initial delay of nearly 30 seconds to a minute and also related to the mechanism of learning and memory retrieval occurring at physiological time-scales has been remaining unsolved.

An alternative question that can be raised is "Are there possibilities for the generation of additional routes through which potentials can arrive at the recording CA1 neuron?" Are all the spines at the localized area between the electrodes readily LINKable with the neighboring ones? If they are not, how can it influence the outcome of the experiment? Can their formation be explained by time-consuming events for the formation of sufficient number of IPLs to achieve sufficient electrical connections towards the recording CA1 neuron? IPL formation necessitates spatial proximity of the converging associated synaptic inputs (Hardie and Spruston, 2009) and also matches with the convergence of inputs for the associative property of LTP (Levy and Steward, 1979). If islets of inter-LINKed spines are formed during LTP induction that also get inter-LINKed with the spines of the recording CA1 neuron, then potentials can reach the recording electrode through multiple routes within a large islet. This can provide an explanation for the potentiated effect of LTP (Fig. 5). Suitability of IPLs in explaining various findings is explained in the following sections.

**Figure 5.**
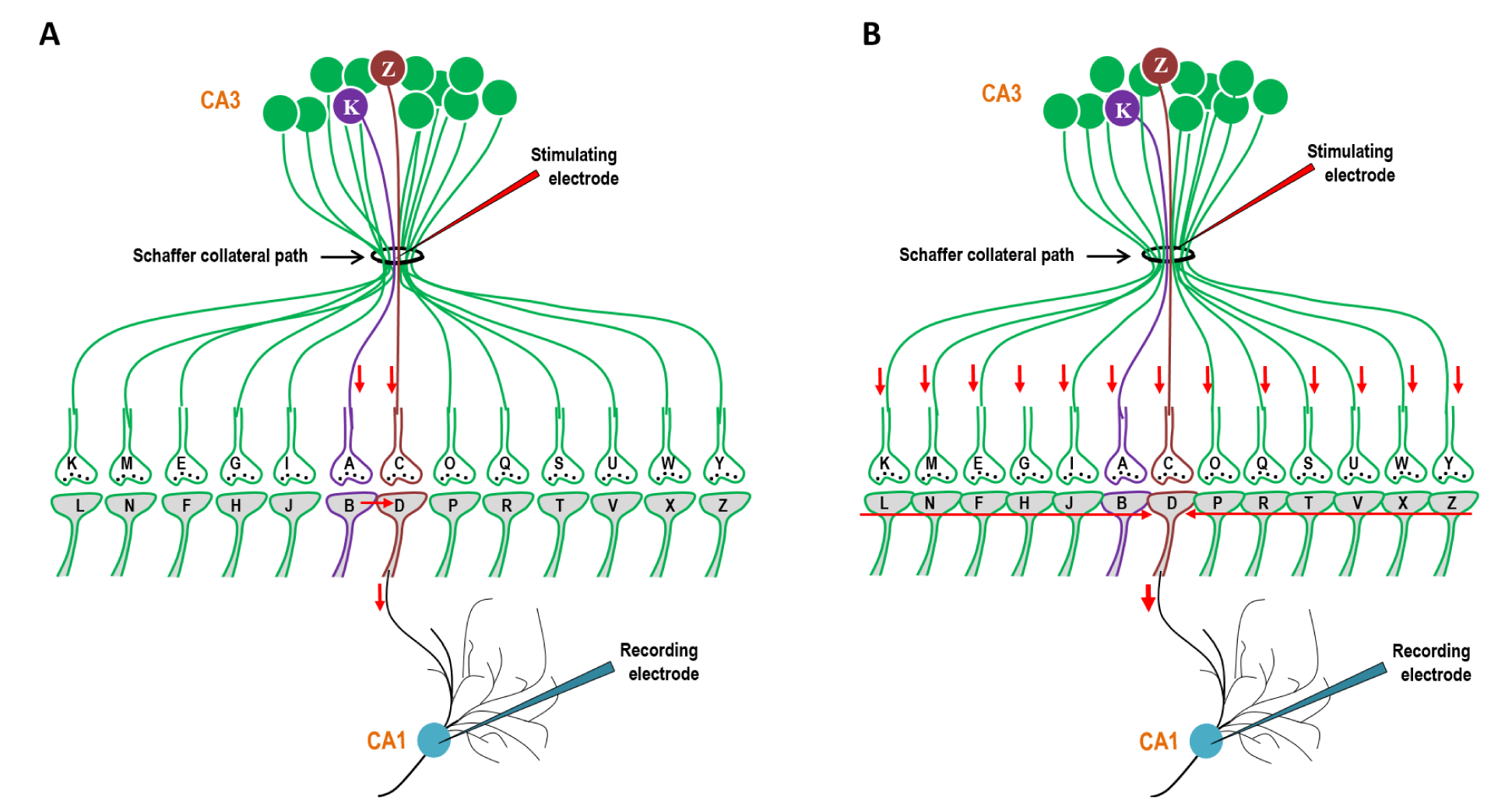
Comparison showing the effect of stimulating the Schaffer collaterals by a regular stimulus before and after inducing LTP. A) Stimulation with a regular stimulus at the Schaffer collateral path before LTP induction will only allow the potentials to propagate through the existing IPL labelled B-D that will propagate through postsynaptic terminal D to its CA1 neuron from which recording is carried out. Even though all the synapses are activated, the EPSPs from postsynaptic terminals B and D only reach the recording CA1 neuron. B) The effect of a regular stimulus at the Schaffer collateral path after the LTP induction will depend on different changes induced by the high-energy stimulation of LTP. The time-delayed formation of different types of IPLs produces increased electrical continuity between the stimulating and recording electrodes. This will allow more current to propagate through all the newly formed IPLs L-N-F-H-J-B and D-P-R-T-V-X-Z(shown by two red horizontal arrows pointing to each other) and reach towards the recording CA1 neuronal soma, which explains the potentiated effect. Note that neurons K and Z are CA3 neurons through which inputs arrived during associative learning (see figure 4).

## 3 Evidence for the role of IPLs in LTP induction and learning

### 3.1 Need for high stimulation energy

High-energy stimulation is necessary for LTP induction. During stimulation, most IPLs are expected to form by close contact between the postsynaptic terminals by excluding water of hydration, a process that requires large amount of energy as evident from studies using lipid membranes (Rand and Parsegian, 1984; Leikin et al., 1987; Cohen and Melikyan, 2004). Additional energy will be necessary for the formation of hemifusion and fusion events as evident by the need for fusogenic molecules to achieve fusion in *in vitro* assays (Keidel et al., 2016). In contrast to the physiological conditions where most IPLs are expected to form only between readily LINKable spines, the high energy released during LTP induction is required for the time-consuming enlargement of the small spines so that they can get abutted to the neighboring spines and undergo IPL formation. An optimal number of IPLs is required for the formation of an optimum number of routes for the regular stimulus-induced potentials to arrive at the recording neuron and show potentiated effect in LTP experiments.

### 3.2 Postsynaptic terminal is the final common path for LTP induction

In experiments conducted after blocking the N-methyl-D-aspartate receptor (NMDA) receptors, a rise in the postsynaptic Ca^2+^ via voltage-sensitive calcium channels produces a potentiated effect (Grover and Teyler, 1990; Kullmann et al., 1992). This shows that the final common change that leads to the potentiated effect occurs at the dendritic spines. This matches with the hypothesized formation of IPLs between the spines as the basic cellular mechanism during LTP induction and correlated changes during learning. Since the aim of the above experiments was to find the final synaptic locations involved during LTP induction, they were conducted by blocking synaptic transmission. However, normal synaptic transmission will be necessary for IPL formation at physiological conditions.

### 3.3 Delay in the induction of LTP

A time delay of 30 seconds (Gustafsson and Wigstrm, 1990) and even more than a minute is observed before the recorded potentials reach the peak level after LTP stimulation. It is also seen (an interpretation from the graphs) in experiments where a transient potentiated effect was produced by a rise in the postsynaptic Ca^2+^ after blocking the NMDA receptors (Grover and Teyler, 1990; Kullmann et al., 1992), single spine LTP experiments (Matsuzaki et al., 2004) and in LTP induction by a single burst (Remy and Spruston, 2007). The tetanus-induced rise in the postsynaptic [Ca^2+^] lasting at most for 2-2.5 s was found sufficient to generate LTP (Malenka et al., 1992). The remaining time delay is not due to the emergence of filopodia or new spines as they take at least 20 minutes to develop (Engert and Bonhoeffer, 1999; Maletic-Savatic et al., 1999). It is also not due to multiple spine synapses between a single axon terminal and a dendrite, as it takes a similar time delay as above (Toni et al., 1999). These findings necessitate the existence of a reversible cellular mechanism that can explain potentiation of up to 300% following a time delay after the induction of LTP and can also explain the correlation with the change occurring during associative learning.

Can the formation of IPLs and islets of inter-LINKed spines explain the time delay between the LTP stimulation and the peak potentiated effect? Are there any similar delays reported in cell biological experiments? Experiments of membrane fusion between cells under the influence of electrical stimulation (Zimmermann and Vienken, 1982; Neil and Zimmermann, 1993; Rems et al., 2013) show comparable delays of minutes. Delays of similar time-scales have been observed for achieving membrane hemifusion (Xu et al., 2005) and fusion (Hofmann et al., 2006; Brunger et al., 2015) by using different SNARE protein properties. Since the average distance between the stimulating and reocording electrodes during LTP experiments is nearly 500*µ*m and since the diameter of the spine heads are nearly 400nm, formation of nearly a thousand IPLs between the spines of different CA1 neurons will be necessary to make electrical continuity through the area between the stimulating and recording electrodes. Since *in vitro* cell fusion is a slow process, the nearly 30 seconds (Gustafsson and Wigstrm, 1990) or even more than a minute of delay to obtain maximum connectivity through the CA3-CA1 synaptic area between the stimulating and recording electrodes can explain the delay during LTP induction (Fig. 6). In this context, the observations that LTP is associated with the enlargement of spine heads (Lang et al., 2004; Matsuzaki et al., 2004) explain the suitability of spine enlargement for the formation of IPLs during the delay period of LTP induction.

**Figure 6.**
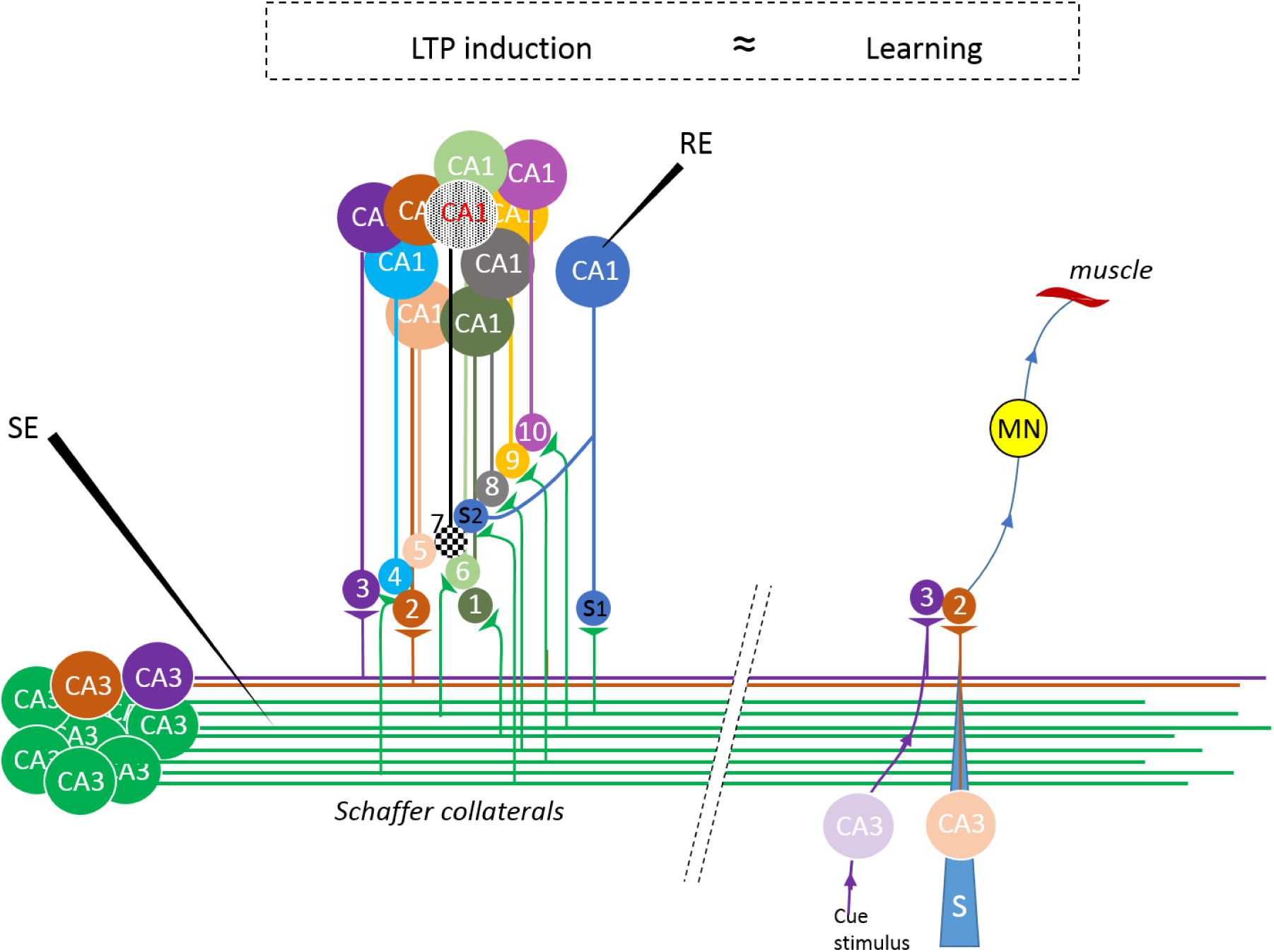
Formation of inter-postsynaptic LINKs (IPLs) that lead to the findings of LTP and its correlation with learning. Left side: A group of CA3 neurons and their recurrent Schaffer collaterals are shown. The recorded CA1 neuron has only two spines that synapse with axonal terminals from stimulating electrode (SE). After stimulation, LTP induction requires arrival of increased current at the recording CA1 neuronal soma after a delay, in response to a regular stimulus at the SE. Several spines (marked from 1 to 5 and 8 to 10 that belong to different CA1 neurons) are activated by the SE. High-energy stimulus used as LTP stimulus lead to slow enlargement of the spines 1 to 5 and 8 to 10. This leads to the formation of different types of IPLs. Spine 7 (checkered spine) is located between the spines 5, 6 and S2. When the latter spines enlarge, they get abutted with the spine 7 and form IPLs with it. These events increase the electrical connection between the SE and RE by many-folds and is only limited by the current carrying capacity of spine S2 and its spine neck. The formation of large number of IPLs between spines including those with interposed non-stimulated spines is a time-requiring process as evidenced from cell fusion studies. Note that nearly a thousand spines are expected to be involved in the formation of IPLs in the volume (IPLs span within the three-dimensional space) between the electrodes. Right side: Formation of a learning-induced IPL between readily LINKable spines marked 3 and 2. Following the learning, the arrival of a cue stimulus at spine 3 reactivates IPL between spines 3 and 2 and induces semblance (S) for memory. The synapses are inverted compared to that of Fig.2. The potentials from inter-LINKed spine 2 can trigger the action potential of a sub-threshold activated motor neuron (MN) that leads to the motor action.

Not all the spines are abutted to each other within the defined space between the stimulating and recording electrodes. Therefore, it will take time for their enlargement to become inter-LINKable for the IPL formation. During the delay for LTP induction, exocytosis of GluR1 receptor subunits of *α*-amino-3-hydroxy-5-methyl-4-isoxazolepropionic acid (AMPA) receptors is expected to take place and facilitate the formation of IPLs between the spines (see section 2.7) for achieving electrical continuity between the electrodes. In contrast during associative learning, it is the readiness of the abutted spines to form IPLs at physiological time-scales that favor learning-induced changes. This explains that IPL formation is a common shared mechanism present during learning and LTP induction.

### 3.4 Sudden drop in the peak-potentiated effect

LTP recordings show that after reaching the peak, the potentiated effect suddenly decreases. The short lasting component of the potentiation is called short-term potentiation (STP). IPL formation by close contact between the postsynaptic membranes, by excluding the water of hydration that requires large amount of energy, is the most common type of IPL. Due to the lack of continuous energy supply, several of these IPLs are expected to reverse back rapidly. This leads to a sudden loss of electrical continuity between the spines of different CA1 neurons that leads to a sudden decrease in the potentiated effect at the recording electrode. This explains STP. Since the qualia of inner sensations of during the retrieval of working, short- and long-term memories are identical and concurrent behavioral motor activities can be produced during the retrieval of these different memories, reactivation of the short-lived IPLs formed by exclusion of water of hydration provides a suitable explanation for working memory. Formation of IPLs by exclusion of water of hydration between the spines and their reversal is expected to occur continuously as the animal interacts with the changing environment. Depending on different factors such as membrane composition and extracellular matrix (ECM) properties, different regions of the nervous system can exhibit different levels of STP.

### 3.5 Correlation between behavioral markers of memory retrieval and LTP

How do the behavioral motor actions indicative of memory retrieval in response to a cue stimulus and the potentiated response to a regular stimulus that demonstrate LTP correlate? The potentials induced by the cue stimulus reactivate the IPLs and activate the inter-LINKed spine to induce inner sensations. The potentials from the inter-LINKed spine propagate towards its soma that lead to motor action (Vadakkan, 2013) (Fig. 4). Since the qualia of inner sensations of retrieved working, short- and long-term memories are identical and concurrent behavioral motor activities can be produced during the retrieval of these different memories, differences in the mechanisms of IPL formation that determine the duration of their maintenance provide a suitable mechanism for different types of memories. The readiness for the formation of IPLs determines the ability to learn and the propensity to form IPLs at a location determines the strength of LTP at that location.

### 3.6 Role of synaptic transmission during learning and LTP induction

From Fig. 4, it can be seen that synaptic transmission at the synapses activated by the associatively-learned stimuli is essential for IPL formation. Similarly, synaptic transmission at the synapses through which the cue stimulus arrives is essential for activating the inter-LINKed spine to induce units of inner sensation. The finding that blockers of the NMDA glutamate receptors block LTP (Collingridge et al., 1983; Herron et al., 1986) supports the observation that normal glutatmatergic synaptic transmission and activation of NMDA receptors lead to the induction of LTP at the location between the CA3 and CA1 neuronal orders at specific stimulating conditions (Morris et al., 1986). Since synaptic transmission is essential for the formation and reactivation of the IPLs, normal synaptic transmission is essential for learning and memory retrieval. It is also necessary for LTP induction and demonstration of LTP maintenance by application of stimuli from the presynaptic side.

### 3.7 Stimulation intensity determines the need for AMPA receptor exocytosis

Even though there is consensus that LTP is mediated by the synaptic insertion of GluR1-containing receptors (Granger et al., 2013), a near-saturation LTP induction alone is sufficient to induce LTP without requiring GluR1 AMPA receptor subunits, their C-tails, or their auxiliary subunits (Herring and Nicoll, 2016). This indicates that membrane interactions between the dendritic spines is the primary cellular mechanism for LTP as explained by the IPL mechanism. At physiological conditions, the readily available GluR1 subunit vesicles are expected to contribute to the IPL formation during learning at physiological time-scales. The observation of GluR1 receptor subunits on the postsynaptic membrane 25nm away from the synaptic junction in normal conditions (Jacob and Weinberg, 2015) indicates the suitability of the lateral regions of the postsynaptic membrane for utilizing the vesicle membrane segments for IPL formation and GluR1 subunits for assembling into functional AMPA receptors.

It is expected that several biochemical and cellular events will continue to take place during the delay period following LTP stimulation to LTP induction. Therefore, these observations can be viewed as a continuum of the changes taking place during normal learning and can be viewed as homeostatic changes that prepares the cell for continued learning. It was found that the GluR1 AMPA receptor subunits redistribute into the cytoplasmic volume of the spine head region after the induction of LTP (Shi et al., 1999; Passafaro et al., 2001). Investigations showed that the spine geometry is critical for AMPA receptor expression (Matsuzaki et al., 2001). Later, it was found that the tetanic stimuli that induce LTP lead to both AMPA receptor insertion and generalized recycling of membrane from endosomes that contain GluR1 AMPAR subunits (Park et al., 2004). Further studies showed that exocytosis of the vesicles containing AMPA receptor subunits is associated with their lateral movement during LTP (Makino and Malinow, 2009). These findings suggest reorganization of the lateral regions of the postsynaptic membranes (spines) using the lipid molecules of the membrane segments from the vesicles that carry AMPA receptor subunits during exocytosis. Since it takes nearly 10s for the AMPA receptors to get recruited to the spines in parallel with the increase in spine volume following LTP induction (Patterson et al., 2010), these can be viewed as continuum of the physiological changes taking place following high energy stimulation.

It was shown that the blockade of exocytosis results in severe loss of LTP (Ahmad et al., 2012; Jurado et al., 2013). IPL formation mediated by the AMPA receptor subunit exocytosis can explain how AMPA receptors affect either the threshold or the magnitude of the LTP at sub-maximal stimulation (Herring and Nicoll, 2016). The propagation of potentials through the IPLs can also explain how mEPSP size is increased after LTP induction (Manabe et al., 1992), which is currently thought to occur either by an increase in the number or function of AMPA receptors at the postsynaptic terminals of the recording neuron (Malenka and Nicoll, 1999). LTP is absent at the synaptic area between the CA3 and CA1 regions in the hippocampi when GluR1 subunits are genetically deleted (Zamanillo et al., 1999) and can be contributed by the lack of GluR1 subunit vesicles for postsynaptic membrane reorganization.

### 3.8 Inhibitors of membrane fusion inhibit LTP

An experiment using blockers of the SNARE proteins introduced into the neuronal cytoplasm showed a reduction in LTP (Lledo et al., 1998). Blockers of the soluble NSF (N-ethylmaleimide sensitive fusion protein) attachment protein receptor (SNARE) proteins can access and block any membrane fusion mediated by the SNARE protein. Postsynaptic exocytosis of the GluR1 receptor subunits during LTP requires a unique postsynaptic Q-SNARE protein for vesicle fusion (Jurado et al., 2013). Since the reorganization of the lateral postsynaptic membrane region is expected to occur during exocytosis of the AMPA receptor subunit-containing vesicles and facilitate the formation of different IPLs, blockers of SNARE proteins can inhibit LTP. In comparison, during associative learning only readily available GluR1 subunit vesicles will be contributing to the IPL formation at physiological time-scales. The action of the SNARE proteins for AMPA receptor subunit exocytosis during this period is expected to contribute to the actual physiological mechanism of learning.

Even though SNARE proteins can lead to membrane fusion through the intermediate stage of hemifusion (Xu et al., 2005), the process has the capability to get arrested at the stage of hemifusion through a specific mechanism (Liu et al., 2008). This is supported by the finding that neuronal SNARE proteins have different mechanisms of hemifusion (Xu et al., 2005; Hofmann et al., 2006; Liu et al., 2008). Furthermore, the postsynaptic protein complexin that can block membrane fusion binds to the SNARE proteins and controls the AMPA receptor exocytosis during LTP induction (Ahmad et al., 2012). Thus, the postsynaptic terminal is expected to have an efficient machinery required for IPL formation and its regulation.

### 3.9 Suitability to accommodate non-Hebbian plasticity changes

Different non-Hebbian potentiation changes were reported at the neighbouring regions of the recording CA1 neuron (Schuman and Madison, 1994; Engert and Bonhoeffer, 1997). What cellular mechanism can lead to the arrival of the potentiated effect at the neighbouring CA1 neurons of the CA1 neuron from which recording is carried out? From the knowledge that the final change during LTP induction is taking place through the postsynaptic terminals (Grover and Teyler, 1990; Kullmann et al., 1992) and the spines enlarge during LTP induction (Lang et al., 2004; Matsuzaki et al., 2004), a large number of IPLs are expected to form between the spines of different CA1 neurons located between the stimulating and recording electrodes. After LTP induction, a regular stimulus can propagate through all the inter-LINKed spines and arrive at different CA1 neurons. This can explain the observations of non-Hebbian changes.

### 3.10 Cooperativity, associativity and input specificity

Findings from different stimulation locations using different stimulation intensities were described to exhibit the properties of cooperativity, associativity and input specificity (McNaughton et al., 1978; Levy and Steward, 1979; Andersen et al., 1977) expected of an ideal learning mechanism. These experiments have been carried at the CA3-CA1 synaptic region. An IPL-mediated mechanism can explain these observations as follows.

1. *Cooperativity*: During LTP stimulation, a critical number of presynaptic terminals is expected to be activated in a cooperative manner to provide an intensity-threshold for LTP induction. However, only a small fraction of these presynaptic terminals directly synapse with the CA1 neuron from which recording is carried out. How can activity reach through a fixed number of synapses to the recording CA1 soma and still show highly potentiated effects? Even though it was explained in terms of the need for depolarization to reduce the Mg^2+^ block of the NMDA receptor channels (Mayer et al., 1984; Nowak et al., 1984), the observation of selective increase in non-NMDA component of EPSP during LTP induction (Kauer et al., 1988) indicates that a non-NMDA receptor-involved postsynaptic mechanism is providing the route for arrival of additional potentials to the recording electrode. Such mechanism is also expected during associative learning at physiological time-scales. Since the membrane segments from the vesicles containing AMPA receptor subunits lead to reorganization of the lateral regions of the postsynaptic membrane parallel to the incorporation of functional AMPA receptors to the postsynaptic membrane, IPL formation is a suitable mechanism to explain the above observations. From Fig. 6, it can be seen that for a regular stimulus to arrive at the recording electrode in a potentiated manner, it is necessary to achieve electrical continuity by the formation of a large number of IPLs, resulting in a large islet of inter-LINKed spines that belong to different CA1 neurons occupying the area between the stimulating and recording electrodes (Fig. 7). In summary, a threshold stimulation energy is required to establish a large islet of inter-LINKed spines for generating LTP.
2. *Associativity*: This is explained as the potentiation of a weak input if it is activated at the same time a strong tetanus is applied at a separate but as a converging input. The convergent nature of the inputs allows separate islets of inter-LINKed spines from the weak and strong stimuli to become electrically connected through IPLs that will allow both islets to get connected with that of the recording CA1 neuron. Simultaneous stimulation is important for excluding water of hydration between the neighbouring postsynaptic terminals that belong to different islets, by their enlargement to form different IPLs to make electrical continuity. This will allow a regular stimulus applied at the location of the weak stimulus to traverse through the islets of inter-LINKed spines formed by the strong stimulus, permitting increased current flow towards the recording electrode.
3. *Input specificity*: This property explains that different inputs that are not active at the time of the strong stimulus do not share the potentiation induced by the strong stimulus. Based on the same mechanism that explains associativity, input specificity also depends on the formation of different IPLs that requires their simultaneous activation.

**Figure 7.**
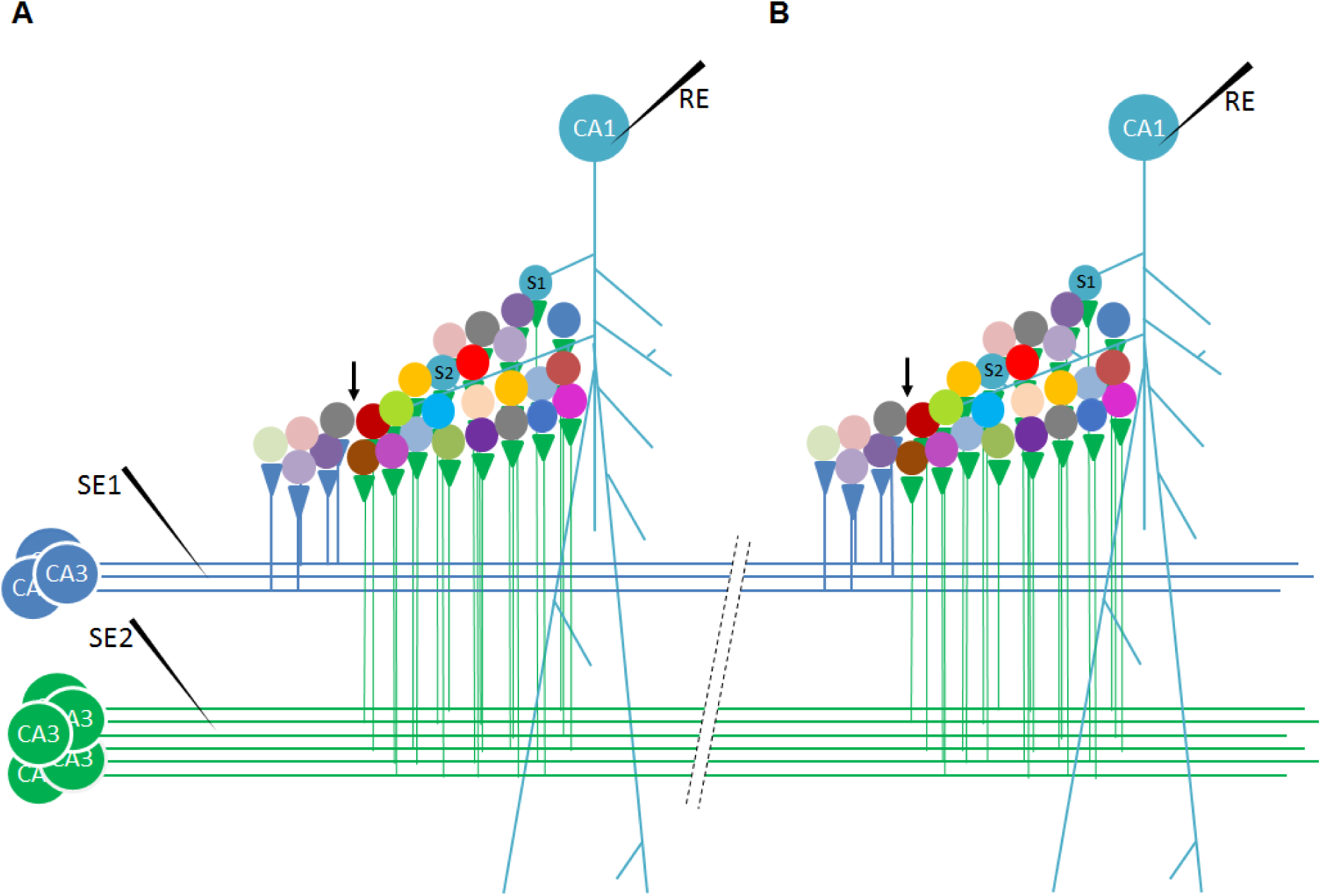
Cellular-level changes of IPL formation that can explain cooperativity, associativity and input specificity. On the left-most side are the CA3 neurons and their Schaffer collaterals that are stimulated by a weak stimulus SE1 through electrode SE1 and a strong stimulus SE2 through electrode SE2. Note that three and six Schaffer collateral axons are activated by the weak and strong stimulus respectively. Also note that the spines are located within a three-dimensional volume (not shown). A) Separate application of weak and strong stimuli provides different results. Weak stimulus SE1 results in the inter-LINKing of spines of five different CA1 neurons (not shown) through IPL formation (shown on the left side of the downward pointing arrow). The formed islet of inter-LINKed spines is not connected with the recording CA1 neuron. Strong stimulus SE2 results in a large islet of inter-LINKed spines that is connected to the recording CA1 neuron. Within the islet, spines that belong to different CA1 neurons inter-LINK with the spines of the recording neuron (S1 and S2). Only a strong stimulus can result in electrical continuity with the CA1 neuron and explains the property of cooperativity. A downward pointing arrow shows that the spines within different islet of inter-LINKed spines formed by weak and strong stimuli remain separated. B) The simultaneous application of weak and strong stimuli can result in the formation of bridging IPLs between the islets of inter-LINKed spines that are formed when they are stimulated separately as in figure A. This is marked by a downward pointing arrow to show that that the formation of bridging IPLs results in electrical continuity between the two different islets of inter-LINKed spines. Following this, the application of a weak stimulus at SE1 will show a potentiated effect when recorded from the CA1 neuron. Simultaneous activation of weak and strong stimuli will result in the removal of water of hydration between the two islets of inter-LINKed spines that are formed by the weak and strong stimuli. This explains associativity. Imagine that another weak stimulus SE3 (not shown) is applied at a different location on the Schaffer collateral. It will share the potentiation induced by the strong stimulus SE2 only if SE3 and SE2 are applied at the same time. Input specificity depends on which weak stimulus is getting simultaneously activated with the strong stimulus S2. From the figure, it is evident that it is necessary to use optimal stimulation strengths and optimal distances between the stimulating electrodes to demonstrate the above properties.

For the demonstration of associativity and input specificity, more than one stimulating electrode need to be used. In order to experimentally demonstrate these features, optimum stimulation intensities have to be applied at optimal distances. This can be explained as follows. Simultaneous activation promotes the formation of critical inter-LINKs between their independently formed islets of inter-LINKed spines to achieve the expected electrical continuity that allows a regular stimulus at the location of application of the weak stimulus to traverse through both the islets of inter-LINKed spines to arrive at the recording electrode. Since islets of inter-LINKed spines can inter-LINK with each other only at specific stimulus intensities and by keeping optimal distances between the stimulating electrodes, experiments to demonstrate input specificity will require optimization of the above parameters. This can also explain the finding that input specificity of LTP is not sustained below a distance of 70m in experiments using hippocampal organotypic slice cultures (Engert and Bonhoeffer, 1997).

### 3.11 Occlusion of learning with LTP

If the same mechanism is responsible for both learning and LTP induction, then large number of spines involved in IPL formation at a specific location by LTP induction can occlude learning that requires spines at that location and vice versa. Both the findings that changes similar to LTP occur during learning (Rogan et al., 1997) and that of occlusion of learning after LTP induction and vice versa (Moser et al., 1998; Whitlock et al., 2006) can be explained in terms of the IPL mechanism. LTP induction leads to a large number of IPLs at a localized area and therefore learning following LTP induction will not to be able to induce any new IPLs at that location. The contribution of the semblance expected to be induced at the localized area, where LTP is induced, towards the net semblance induced by the cue stimulus (from the remaining areas in the hippocampi and the cortices) is likely very small. Therefore, memory should not be reduced for this reason alone. However, the large number of IPLs induced by LTP induction can lead to a large number of non-specific semblances at the time of memory retrieval as the cue stimulus traverses through all these non-specifically inter-LINKed spines. The large non-specific semblance can contribute to the reduced memory observed in these experiments. The single IPLs that are induced sparsely at different locations in the nervous system during learning may not always occlude LTP induced at a localized region. However, using focussed experiments in the hippocampus where a strong convergence of inputs occurs, it was found that learning can occlude LTP induction (Whitlock et al., 2006). Results from occlusion experiments demonstrate that both learning and LTP induction are mediated through the same mechanism and IPL formation is suitable for such mechanism.

### 3.12 Dopamine augments motivation-promoted learning and LTP

Motivation can enhance learning (Wise, 2004). This can be explained by the enlargement of the spines by dopamine (Yagishita et al., 2014) released during motivation-promoted learning, which can augment IPL formation. Since dopamine takes 0.2 to 0.3 seconds for its action (Yagishita et al., 2014), it is likely that motivation is necessary prior to associative learning to have an effect. Enlargement of the spines is likely to increase the duration of maintenance of the IPLs, which in turn can increase the probability for the IPLs to get stabilized. The duration of stabilization of the IPLs in turn can determine the duration of maintenance of associative learning-induced changes. This provides a cellular-level explanation for the motivation-facilitated associative learning. Through a similar mechanism, it also explains how dopamine receptor activation can increase the magnitude of LTP in the hippocampal slices (Otmakhova and Lisman, 1996).

### 3.13 Inhibitors of NMDA receptors do not reverse late LTP maintenance

Even though NMDA receptor antagonists block the initial encoding of learning-induced changes in animals, they do not block the maintenance phase of learning-associated changes (Day et al., 2003). In parallel to this, it was observed that inhibitors of the NMDA receptors do not reverse the late maintenance phase of LTP (Ling et al., 2002). This can be explained by the fact that in order to block the learning-induced changes or an already induced LTP, there should be a mechanism to block the functional effect of the formed IPLs. Since inhibition of NMDA receptors cannot have any effect on any type of formed IPLs, they do not reverse either learning-induced changes or an already induced LTP.

### 3.14 Maintenance phase of LTP

A retrieval-efficient learning mechanism can take place within milliseconds that leads to the formation of different type of IPLs. Stabilization of the IPLs can take place if the same learning takes place repeatedly, which is facilitated by the presence of motivation-promoted dopamine release and spine enlargement. The changes occurring following IPL formation replenish the substrates to optimize the structural environment for future learning events. Following LTP induction that generate large number of IPLs, changes similar to that occur following learning-induced IPL formation continue to take place. During both persistence of learning-induced changes and LTP maintenance, different protein molecules of the biochemical cascades will be associated with the maintenance of IPLs. Examples include calcium/calmodulin-dependent protein kinase II (CaMKII) and protein kinase C Mζ (PKC Mζ).

Entry of calcium into the postsynaptic terminal activates the protein CaMKII, which subsequently binds to the NMDA glutamate receptors and phosphorylates principal and auxiliary subunits of AMPA glutamate receptors (Lisman et al., 2012). Since exocytosis of readily available AMPA subunit vesicles occur following LTP induction, it may explain how a) some of the post-LTP induction biochemical intermediates are correlated with associative learning, and b) blocking or genetic deletion of the molecules of these pathways will be blocking LTP induction due to their effect on IPL formation. Inhibitors of PKC Mζ were found to reverse the established LTP (Ling et al., 2002) and lead to the loss of spatially learned information (Pastalkova et al., 2006). The finding that PKC Mζ is concentrated at the location of cell membrane abscission (Saurin et al., 2009) indicates its possible role in regulating the separation of cell membranes, which may prevent reversal of the IPLs. However, the finding that memory and LTP are normal in the PKC Mζ knockout mice (Lee et al., 2013; Volk et al., 2013) indicates that alternate mechanisms will be sufficient to maintain the formed IPLs. After the initial minutes following LTP induction, slow reversal of the inter-postsynaptic membrane fusion and hemifusion are expected to take place along with endocytosis of AMPA receptor vesicles (Dong et al., 2015).

### 3.15 LTP: Kindling: Memory: Seizure

Kindling is induced by stronger stimulation energy than those used for LTP induction and can cause afterdischarges. Kindling shows several similarities to human seizure disorders (Bertram, 2007). This can be explained in terms of the conversion of the IPL mechanism of hemifusion to fusion (Vadakkan, 2016b) and provides a suitable comparable change occurring *in vivo*. The fused areas allow the propagation of potentials between their postsynaptic terminals without any additional resistance and remain for long period of time. This can explain the findings in kindling experiments. Fusion changes can explain the transfer of injected dye from one CA1 neuron to the neighboring ones observed in animal models of seizures (Colling et al., 1996). It has been observed that the gene expression pro les of adjacent CA1 are different (Kamme et al., 2003; Cembrowski et al., 2016). Therefore, mixing of the cytoplasmic contents, though the membrane fusion, even between similar types of neurons that have different gene expression pro les can lead to the triggering of homeostatic mechanisms such as spine loss as seen both after kindling (Singh et al., 2013) and in seizure disorders (Swann et al., 2000).

### 3.16 Forgetting and reversal of LTP

The potentiated effect caused by raising the postsynaptic Ca^2+^ via voltage-sensitive calcium channels at the glutamatergic synapses (Kullmann et al., 1992) and the potentiation of AMPA currents by single spine LTP experiments (Matsuzaki et al., 2004) last only for nearly 30 min. This indicates that the formed IPLs reverse back quickly. Since most IPLs are formed by close contact between the spine membranes by exclusion of the water of hydration, which requires a large amount of energy, they reverse back quickly explaining the reversal of the potentiated effect within a short period of time. This can provide a mechanism for working memory. Following the initial phase of rapid reversal responsible for STP, LTP reverse back very slowly. This slowly reversing phase has similarities to the slow reversal of changes for those memories that last beyond working memory. It can be explained in terms of the formation, stabilization and reversal of inter-spine membrane hemifusion. Slow reversal of these changes is expected to associate with endocytosis of the AMPA receptor subunits and reduction in the size of enlarged spines leading to LTP decay (Dong et al., 2015).

### 3.17 Testing of LTP by regular stimulus may maintain the IPLs

After LTP induction, any ordinary stimulus applied at the stimulating electrode traverses through the large number of newly formed IPLs and reaches the recording electrode patch-clamped to the CA1 soma, which is recorded as a potentiated effect. Each of these regular stimuli at frequent intervals reactivates the already formed IPLs and may even have a role in maintaining them, which can be verified. Repeated reactivation of the existing IPLs by the regular stimulus for recording the potentiated effect is similar to the repeated retrieval of memories.

### 3.18 LTP and place cell firing

Role of IPLs in triggering place cell firing was explained previously (Vadakkan, 2016a). LTP induction is known to modify specific sets of place cells (Dragoi et al., 2003) indicating that the formation of a large number of new IPLs induced by LTP can lead to the spread of potentials through these IPLs and result in the firing of additional postsynaptic CA1 neurons that are being held at a sub-threshold state. A similar mechanism can explain the incremental re-mapping of the CA1 place cells to a final fully-differentiated form following environmental experience (Lever et al., 2002). In both LTP induction and learning, new IPLs are formed, which can provide additional potentials to those neurons that are at a sub-threshold activated state under baseline conditions (Fig. 4).

### 3.19 Blocking the extracellular matrix space blocks LTP

Based on the explanations provided by the IPL-mediated mechanisms, blocking the ECM space between the spines can prevent IPL formation and LTP induction. Perineural net proteins around the spines of the CA2 region of the hippocampus can explain the reduced LTP in this region and improvement in LTP following removal of those proteins (Carstens et al., 2016). Since any resistance to the IPL formation is expected to prevent their rapid formation in chains (Vadakkan, 2016b), resistance offered by the perineural net proteins to IPL formation can support the finding that CA2 region is uniquely resistant to seizures (Hatanpaa et al., 2014).

### 3.20 Dendritic spikes and LTP

The proposal of dendritic spikes as a mechanism for co-operative LTP (Golding et al., 2002) and the findings that dendritic spikes are necessary for single-burst LTP (Remy and Spruston, 2007), and that Ca^2+^ spikes cause long-lasting potentiation of spines active at the time of spike generation (Cichon and Gan 2015) can be explained on the basis of large number of IPLs formed between the spines of different neurons. The formation of a large islet of inter-LINKed spines can explain the observation of single-burst LTP. This matches with the explanation that a dendritic NMDA spike is a synchronous activation of 10 to 50 neighboring glutamatergic synapses triggering a local regenerative potential (Antic et al., 2010) and can explain the functional significance of dendritic spikes *in vivo* (Sheffield and Dombeck, 2015). The shared physical properties of the environment in which the nervous system operates are likely responsible for the large inter-LINKed spines and the semblances induced by the dendritic spikes are likely contributing to the background semblance of the system. The lateral spread of activity through the inter-LINKed spines during NMDA spikes is expected to have a significant contribution to one of the vector components of the oscillating extracellular potentials at rest. If there are a large number of stabilized IPLs present at the distal dendritic compartment inter-LINKing several spines (as evident from dendritic spikes), then based on the present work the requirement for AMPA receptor subunit vesicle exocytosis and spine membrane reorganization at these locations will be less. A lesser number of AMPA receptors at the distal areas of the dendritic tree compared to the proximal area at the stratum lacunosum-moleculare region of the CA1 pyramidal neurons (Nicholson et al., 2002) supports such an expectation.

### 3.21 Single spine LTP

A paired stimulation protocol by uncaging glutamate has allowed the stimulation of individual spines of CA1 neurons (Matsuzaki et al., 2004). Recording from the soma of the neuron showed potentiation of AMPA-receptor mediated currents (AMPA currents). In this experiment, the potentiation effect occurred following a delay of 3 minutes after the stimulation and lasted for nearly 30 minutes. The potentiation was observed only when the spine had undergone enlargement. Potentiation of AMPA currents at the small spines was strongly correlated with the enlargement of these spines, indicating that the latter has contributed to the formation of IPLs with abutted spines (that belong to other CA1 neurons). Since recording was carried out from the CA1 soma, IPL formation with the spines of other CA1 neurons can be expected to result in the arrival of AMPA currents from the latter. The magnitude of the potentiation was also correlated with early or long-lasting spine enlargement indicating the formation of a maximum number of IPLs with the abutted spines, which can increase the current flow towards the recording electrode through the newly formed IPLs. In contrast, the spines that are already large at the baseline state are likely to have existing IPLs and lack any free surface area to form additional IPLs. This may explain why they did not show much potentiated effect. The time delay of 3 minutes can explain the time required for spine expansion and IPL formation as explained in section 3.3. Since high-energy is required to bring the spines to close contact with each other, they can reverse reverse back relatively quickly and can explain why the duration of potentiated effect is limited to only 30 minutes in these experiments.

## 4 Predictions

1. A spectrum of IPL changes are expected to take place during LTP induction. This ranges from exclusion of water of hydration between the spines to fusion, with intermediate stages of partial and complete hemifusions, between the abutted spines of the CA1 neurons.
2. The strength of LTP induced at different locations of the nervous system for a given distance between the stimulating and recording electrodes depends on the number of IPLs formed and inter-LINKed with each other during the delay time after stimulation capable of conducting current to the recording neuronal soma. This in turn depends on the number of converging inputs, their density, inter-spine ECM properties and lipid composition of the postsynaptic membranes at different locations.
3. Kindling induced at the Schaffer collateral area is expected to produce more inter-spine fusion than by LTP induction at the same location. Moreover, the area of inter-spine fusion is expected to be larger than that is induced by LTP. Those large fused areas are expected to be irreversible when compared to mostly small and reversible ones that are likely induced by LTP.

## 5 Discussion

The arrival of increased current at the recording electrode following high energy stimulation at the Schaffer collaterals has been examined in the context of the Hebbian proposal that strength of individual synapses change during learning. As a result, the above observation has been viewed as resulting from increase in the strength of the individual CA3-CA1 synapses. Even though LTP studies based on the Hebbian view have provided several features for model fitting, investigators have been expecting to find a perfect match (Bliss and Collingridge, 1993; Malenka, 2003; Nicoll,2017). What remains unmatched in the matching process using Hebbian view include a) the delay of at least 30 seconds for LTP induction in terms of physiological time scales of milliseconds at which learning-induced changes and memory retrieval take place, b) the delayed biochemical changes observed after LTP induction and their correlations with normal learning and LTP, and c) how LTP is correlated with working memory. Therefore, it was necessary to find an alternate mechanism for explaining all the findings in an inter-connectable manner.

The term potentiation was used in describing the finding of the arrival of increased current at the recording electrode in response to a regular stimulus by assuming that the changes are occurring at a fixed number of synapses at the dendritic spines of the recording CA1 neuron. Since explanations for almost all the findings associated with LTP have become possible by the IPL-mediated mechanism and since it involves nearly a thousand dendritic spines, LTP can be explained as resulting from a formation of a chain of IPLs at the CA3-CA1 synaptic area. The predictions made by the derived mechanism are testable.

The potentials arriving through the IPLs also provide efficient mechanistic and functional explanation for LTP that previously thought as a) enhancement of neurotransmitter release (Malinow and Tsien, 1990), b) increased postsynaptic sensitivity to glutamate (Manabe and Nicoll, 1994) and also as an expression-mechanism (Nicoll and Malenka, 1999) taking place at a fixed number of synapses with the spines of the recording CA1 neuron. It was found that following LTP stimulation, functional AMPA receptors get inserted to the postsynaptic membranes of silent synapses that do not have any functional AMPA receptors and convert them to functional synapses (Isaac et al., 1995; Liao et al., 1995). Readily available vesicles containing AMPA receptor subunits during learning can provide both its lipid membrane segments for reorganization of the lateral aspects of the spines for IPL formation and GluR1 subunits for the formation of functional AMPA receptors. Same changes are expected to occur for a prolonged period after LTP stimulation.

The different types of IPLs provide much-needed flexibility to explain the retention of learning-induced changes for different durations to explain memories that can be retrieved after corresponding durations. This matches with the expectation of a single mechanism expressed ubiquitously throughout the nervous system for different types of memories (Shors and Matzel, 1997). IPL mechanism provides a universal effect for the action of different neurotransmitters. For example, activation of muscarinic acetylcholine receptors leading to a potentiated effect at the recording CA1 pyramidal neuron (Dennis et al., 2016) can be explained in terms of IPL formation. Effect of IPLs between closely located spines can be viewed from different electrophysiological findings. For example, as new granule neuron dendrites branch out towards the performant path axonal terminals, they form IPLs with the pre-existing islets of inter-LINKed spines. This is observed either as an increase in the amplitude or a reduction in threshold for inducing LTP (Schmidt-Hieber et al., 2004; Ge et al., 2007).

Formation of islets of inter-LINKed spines necessitates related learning events to share several IPLs, making learning more efficient. Schemas of previous associatively-learned items or events induced as semblances at the islets of inter-LINKed spines can be used during related learning events (Tse et al., 2007). It also allows avoidance of concerns about saturation of synapses or overwriting of connections (Bliss, 1998; Fusi and Abbott, 2007). In contrast to the clustered plasticity model (Govindarajan et al., 2006), the islets of inter-LINKed spines that belong to different neurons (exceptions can be present) provide a structure-function explanation for the generation of internal sensations together with the mechanism for stimulus-specific motor activation. The fast mechanisms among the spectrum of IPL changes enable learning at physiological time scales and allow instantaneous retrieval of memory following the learning without having to wait for the occurrence of biochemical sequence of events. This can bypass the need for searching for a separate mechanism for working memory. In summary, the present work has demonstrated a cellular mechanism that can correlate learning and LTP induction and provides an opportunity for crossing the explanatory chasm between different fields of investigations of the nervous system.

## Acknowledgements

This work was supported by Neurosearch Center, Toronto. I would like to thank Selena Beckman-Harned for reading the manuscript.

### Conflict of interest

U.S. patent: number 9477924 pertains to an electronic circuit model of the inter-postsynaptic functional LINK.

